# *De novo* mutations in regulatory elements cause neurodevelopmental disorders

**DOI:** 10.1101/112896

**Authors:** Patrick J. Short, Jeremy F. McRae, Giuseppe Gallone, Alejandro Sifrim, Hyejung Won, Daniel H. Geschwind, Caroline F. Wright, Helen V. Firth, David R. FitzPatrick, Jeffrey C. Barrett, Matthew E. Hurles, on behalf of the DDD study

## Abstract

**Summary:** *De novo* mutations in hundreds of different genes collectively cause 25-42% of severe developmental disorders (DD). The cause in the remaining cases is largely unknown. The role of *de novo* mutations in regulatory elements affecting known DD associated genes or other genes is essentially unexplored. We identified *de novo* mutations in three classes of putative regulatory elements in almost 8,000 DD patients. Here we show that *de novo* mutations in highly conserved fetal-brain active elements are significantly and specifically enriched in neurodevelopmental disorders. We identified a significant two-fold enrichment of recurrently mutated elements. We estimate that, genome-wide, *de novo* mutations in fetaLbrain active elements are likely to be causal for 1-3% of patients without a diagnostic coding variant and that only a small fraction (<2%) of *de novo* mutations in these elements are pathogenic. Our findings represent a robust estimate of the contribution of *de novo* mutations in regulatory elements to this genetically heterogeneous set of disorders, and emphasise the importance of combining functional and evolutionary evidence to delineate regulatory causes of genetic disorders.

## Main Text

The importance of non-coding variation in complex disease is well established–the majority of disease-associated common SNPs lie in intergenic or intronic regions, albeit with low effect sizes^1,2^. Rare sequence and structural variants in relatively few regulatory elements have been causally linked to specific Mendelian disorders^3,4^ (reviewed in *Mathelier et. al, 2015* and *Spielmann and Mundlos, 2016*), most of which act dominantly, with a few exceptions^5^. These pathogenic regulatory variants can act by loss-of-function ^6–9^ or gain-of-function ^10,11^. These regulatory elements can lie far from the gene they regulate. For example, a set of sequence variants in an evolutionarily conserved regulatory element located 1Mb from its target gene, *SHH*, can cause polydactyly^10^. As a consequence, in both complex and Mendelian disorders it can often be challenging to identify the gene whose regulation is being perturbed by an associated regulatory variant^12–14^. Moreover, the contribution of highly penetrant mutations in regulatory elements to highly genetically heterogeneous rare diseases, such as neurodevelopmental disorders, has not been firmly established.

We recruited 7,930 individuals with a severe, undiagnosed developmental disorder, and their parents to the Deciphering Developmental Disorders (DDD) study from clinical genetics centres in the UK and Ireland. Systematic clinical phenotyping^15^ identified 79% with intellectual disability (87% with more broadly defined neurodevelopmental disorders). Congenital heart defects were the most prevalent non-neurodevelopmental phenotype recorded, present in 10% of the cohort. Exome sequencing of these families revealed that damaging *de novo* mutations (DNMs) in known DD-associated genes are present in ~25% of probands, accounting for the majority of diagnostic variants^16,17^, and that an additional 17% of probands carry pathogenic DNMs in genes not yet robustly associated with DDs^17^. Thus the majority of the probands do not carry a diagnostic variant in a protein-coding gene, and are termed ‘exome-negative’. To explore the role of DNMs in non-coding, likely regulatory, elements we performed targeted sequencing on three classes of putative regulatory elements: the 4,307 most evolutionarily conserved non-coding elements (CNEs)^18^, 595 experimentally-validated enhancers^19^, and 1,237 putative heart enhancers^20^, together covering 4.2Mb of sequence. Furthermore, we define a set of ‘control’ non-coding elements covering 6.03Mb by analysing intronic sites at least 10bp away from the nearest exon that are targeted by exome baits and were sequenced to >30X depth (see Methods).

## Selective constraint acting on non-coding elements

We first assessed how much purifying selection had skewed allele frequencies in non-coding elements. We used the mutability-adjusted proportion of singletons (MAPS) metric^21^ in 7,080 unrelated, unaffected DDD parents to test six different element classes: introns, heart enhancers, validated enhancers, CNEs, protein-coding genes, and known DD-associated genes. Both the introns and the heart enhancers, like synonymous variants in proteinucoding genes, show little evidence of purifying selection (Figure 1a). The experimentally-validated enhancers and CNEs are constrained to a similar degree to all protein-coding genes, but less than known DD-associated genes (Figure 1a). Statistical power to detect functionally relevant variants in protein-coding genes is strengthened considerably by stratification of variants on the basis of their likely impact on the encoded protein (e.g. missense and truncating variants) and using metrics that predict variant deleteriousness such as CADD^22^. We used the MAPS framework to assess whether deleteriousness metrics were predictive of selective constraint in the targeted non-coding elements. We computed the MAPS within bins of CADD scores encompassing 1,520,250 variants in unaffected DDD parents. In protein-coding genes, the strong correlation between CADD score and strength of purifying selection enabled differentiation between variants that are neutral, weakly constrained and very highly constrained. In CNEs and validated enhancers, CADD differentiates neutral variation from those under weak constraint (similar to missense level in the protein-coding regions), but failed to identify highly deleterious variants with selective constraint on a par with protein truncating variants (Figure 1c). Other deleteriousness metrics were assessed, but none were more informative than CADD (Figure S1). These findings replicate across the entire non-coding genome in an independent sample of 3,781 UK whole genomes sequenced to 7x coverage as part of the UK10K project^23^ (Figure S2).

**Figure 1.**
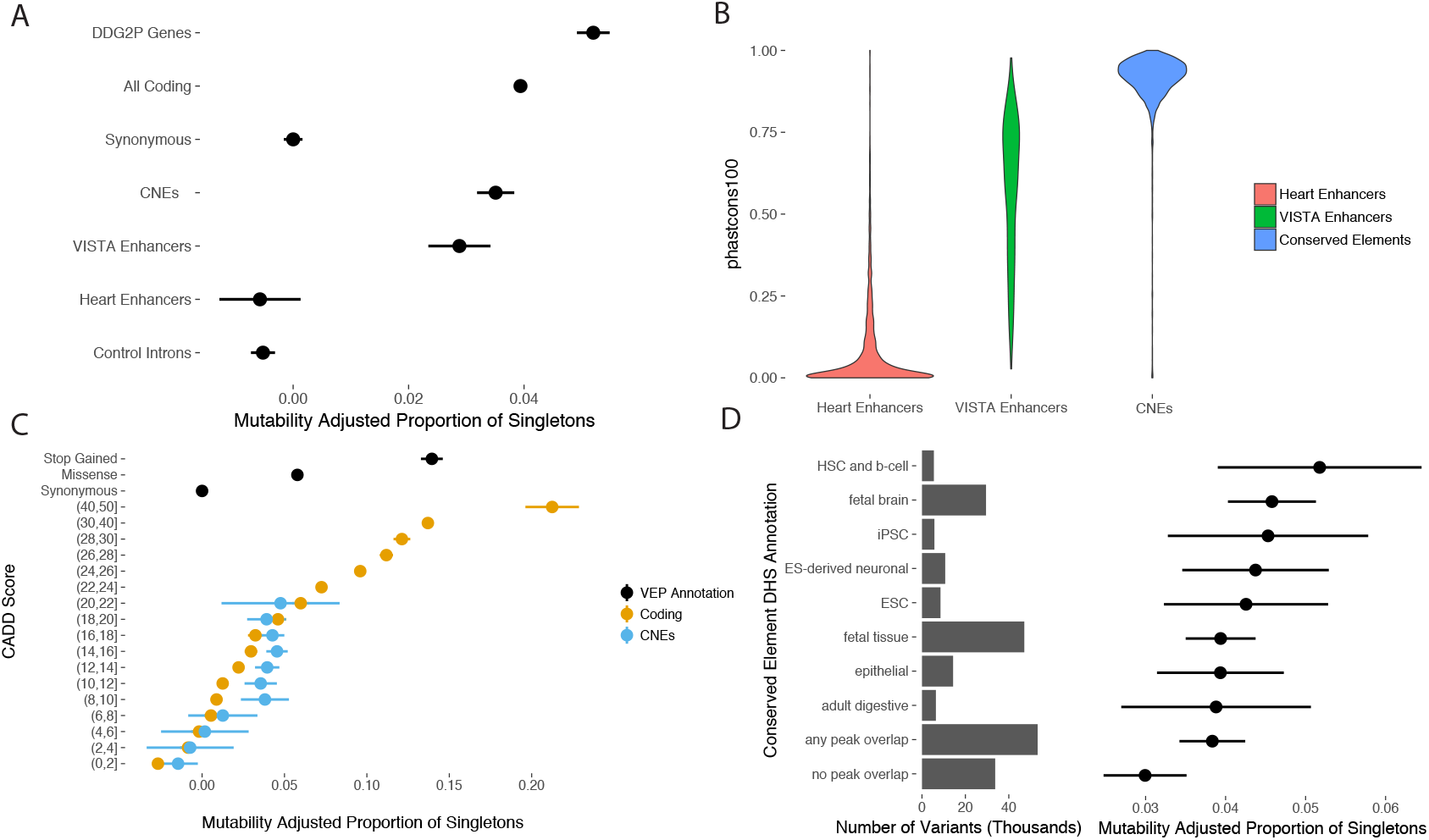
Selective constraint in targeted non-coding elements (a) Strength of selection in targeted non-coding elements compared to protein-coding regions. Introns and putative heart enhancers show little evidence of purifying selection while CNEs show selection on par with all genes, but less than known DD genes. (b) Evolutionary conservation score (phastcons100^18^) for CNEs, experimentally validated enhancers, and putative heart enhancers. (c) Using CADD to stratify coding and non-coding variants observed in unaffected parents. CADD differentiates neutral variation from weakly and strongly constrained sites in coding regions, but fails to identify non-coding variation with selection pressure on par with loss of function variants in coding regions. (d) Purifying selection in regions of open chromatin across different tissues. Sites overlapping a DNase hypersensitive site (DHS) in at least one tissue are under stronger purifying selection than sites not overlapping a DHS.

Functional genomic annotation of putative regulatory elements provides additional information to distinguish elements and sites that are more likely to be functionally relevant for DD-causation. We used DNase I hypersensitivity sites (DHS) in 39 tissues and chromHMM genome segmentation predictions in 111 tissues^24^ to predict tissue activity for the targeted non-coding elements. Of the 4,307 CNEs sequenced in this analysis, 4,046 (93.9%) were active in at least one of the 111 surveyed tissues while 261 (6.1%) were inactive or repressed in all tissues. Most (n=212) of these 261 inactive CNEs exhibited polycom-mediated repression in several tissues suggesting they are active in tissues not yet surveyed. Nonetheless, active CNEs were under stronger selective constraint than inactive CNEs, despite no difference in evolutionary conservation (Figure S3). Likewise, variants within a DHS peak in at least one tissue were under stronger purifying selection than variants not overlapping a DHS peak (p = 0.019) (Figure 1d). Within the set of active elements, we did not find any significant difference in purifying selection in elements active in a single tissue compared to those active in multiple tissues (Figure S3), nor did we identify significant differences in selective constraint in the targeted elements between tissues

## Enrichment of de novo mutations in non-coding elements

We identified candidate *de novo* single nucleotide mutations in 7,930 trios (Methods). We identified 1,691 ‘exome-positive’ individuals with a likely pathogenic protein-altering DNM or inherited variant in a known DD-associated gene, with the remaining 6,239 being ‘exome-negative’. Using a previously described model for germline mutation^25^, we compared the numbers of observed and expected DNMs in the different sets of non-coding elements in exome-positive and exome-negative trios. No significant DNM enrichment was observed in exome-positive probands for any of the sets of non-coding elements, demonstrating that the mutation model is well-calibrated for our data (Figure S4). By contrast, we found that the targeted CNEs (irrespective of their tissue activity) are nominally significantly enriched for DNMs in exome-negative individuals (422 observed, 388 expected, p = 0.04), whereas experimentally-validated enhancers (153 observed, 156 expected, p = 0.596), heart enhancers (86 observed, 86 expected, p = 0.511), and intronic controls (901 observed, 920 expected, p = 0.719) were not enriched (Figure 2). Even when focusing on the minority of probands with heart phenotypes, no enrichment for DNMs in heart enhancers was observed. Subsequent analyses focus exclusively on exome-negative trios.

**Figure 2.**
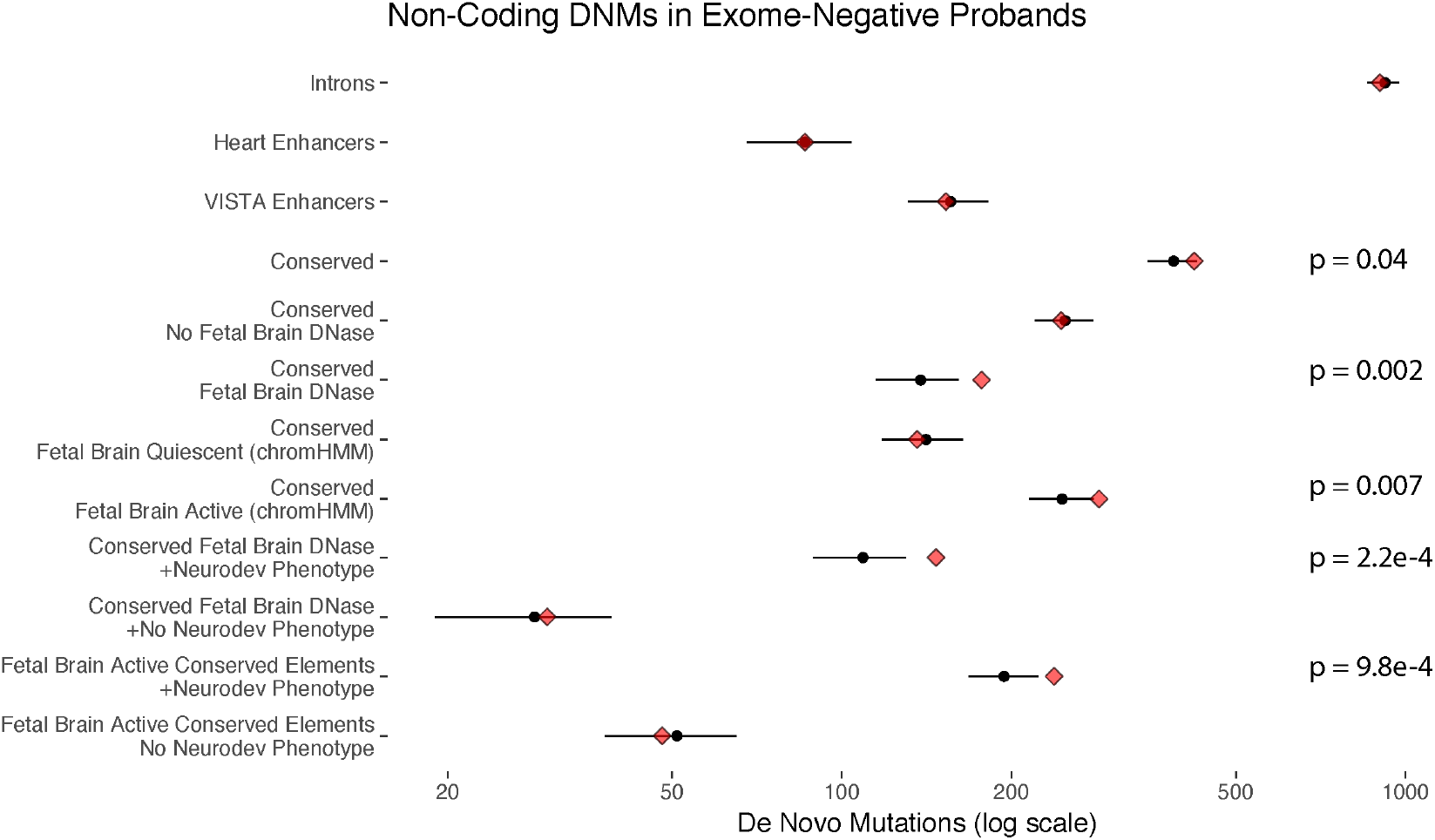
Enrichment of DNMs across targeted non-coding element classes and functional annotations in exome-negative probands. Red diamonds indicate the observed counts, while black circles and bars indicate the expected count and 95% CI. Targeted CNEs showed a modest enrichment for DNMs (422 observed, 388 expected, p = 0.04) while heart enhancers, experimentally-validated enhancers, and control introns matched the null model. The observed enrichment is specific to CNEs predicted to be active in the fetal brain and to patients with neurodevelopmental disorders (238 observed, 194 expected, p = 9.8e-4).

Given the preponderance of individuals with neurodevelopmental disorders in our cohort but the broad range of the tissue activity of the targeted non-coding elements, we refined our testing for DNM enrichment by focusing on CNEs active in fetal brain. We observed a strong significant enrichment of DNMs within 2,077 fetal brain DHS peaks in CNEs (177 observed, 138 expected, p = 0.002) but no enrichment sites in CNEs falling outside of fetal brain DHSs (245 observed, 249 expected, p = 0.623) (Figure 2). We also used chromHMM^26^ predictions of fetal brain activity, which incorporate a broader range of functional assays, and again identified a significant enrichment of DNMs in the 2,613 fetal brain active CNEs (286 observed, 246 expected, p = 0.007) but not in the fetal brain inactive (quiescent, heterochromatin, or polycomb-repressed) elements (136 observed, 141 expected, p = 0.692) (Figure 2). Moreover, the DNMs observed in fetal brain active CNEs in exome-negative probands were at more highly conserved sites (Wilcoxon rank sum test on PhyloP 100-way score^27^) compared to DNMs observed in exome-positive probands (Figure S5). The absence of DNM enrichment in fetal brain inactive elements suggests that modulation of existing enhancer activity in the brain is more likely to cause developmental disorders than gain of ectopic brain activity for elements active in other tissues. The excess of DNMs observed in fetal brain active CNEs and within fetal brain DHS peaks is concentrated exclusively within the 79% of ‘exome-negative’ probands with one or more neurodevelopmental phenotypes (fetal brain DHS peaks: p = 2.2e-4, fetal brain active by chromHMM: 238 observed, 194 expected, p = 9.8e-4), with no significant enrichment observed in those without neurodevelopmental phenotypes (fetal brain DHS: p=0.426; fetal brain active by chromHMM: p=0.671) (Figure 2). In summary, we identified a highly significant and specific enrichment of DNMs in fetal brain active CNEs in exome-negative probands with neurodevelopmental disorders

Having demonstrated a specific enrichment of DNMs in fetal brain active CNEs in probands with neurodevelopmental disorders, we re-evaluated the subset of experimentally-validated enhancers with functional evidence for activity in fetal brain (N=383) in these probands and observed an enrichment for DNMs only within the top quartile of evolutionary conservation (18 observed, 9 expected, p = 0.01 within fetal brain DHS) (Figure S6), suggesting that even for experimentally validated fetal brain enhancers, the DNM enrichment is concentrated within elements with strong evolutionary conservation.

We assessed four different methods for gene target prediction: Genomicus^28^ (based on evolutionary synteny), DHS-expression correlation^29^, Hi-C in fetal brain^30^ and simply choosing the closest gene. The proportion of fetal brain active CNEs for which a prediction could be made varied from 26% (fetal brain Hi-C) to 100% (closest gene). The pairwise-concordance between the four approaches was low: at most ~ 35% between any two methods (Figure S7). We did not identify any enrichment for DNMs in elements predicted to target known DD genes, or likely dosage sensitive genes (defined by the *'*probability of loss of function intolerance*'* (pLI) metric^21^), or genes that are more highly expressed in the brain compared to other tissues (Methods, Figure S7)

We identified 45 transcription factors (TFs) whose motifs were significantly enriched in fetal brain active CNEs (Methods), are expressed in the brain and for which position-weight matrices are available^31^. We assessed the impact of DNMs on predicted TF binding, and observed nominally significant increases in DNMs predicted to increase affinity and overlap of binding sites, but none of these results survived correction for multiple testing (Figure S8). Given the number of DNMs we have identified, and the relative immaturity of *in silico* predictions of the impact of non-coding variation on gene expression, it is not currently possible to determine the mechanisms by which these DNMs contribute to DDs.

To explore the likely penetrance associated with the observed DNM enrichment in the targeted non-coding elements, we investigated potential over-transmission of inherited rare variants in these elements to affected children. No class of non-coding element classes showed any evidence for overtransmission (Figure S9). Furthermore, we did not detect any enrichment for other variants in the same element as the DNM that would suggest the DNM is acting as a second hit to an already *'*perturbed*'* haplotype. These observations are consistent with the enrichment of DNMs in fetal brain active CNEs comprising a mixture of 70-80% non-pathogenic DNMs and 20-30% of highly penetrant dominantly-acting DNMs.

## Recurrently mutated regulatory elements

We observed a significant excess of recurrently mutated fetal brain active CNEs (two or more DNMs in unrelated individuals) compared to the expectation under the null mutation model (31 observed, 15 expected, p = 1.2e-4) (Figure 3a). However, no individual CNE exceeds a conservative genome-wide significance threshold of p<1.91e-5 (Bonferroni-corrected p-value based on independent tests for enrichment across 2,613 fetal brain active elements) (Figure 3b). We did not observe any significant clustering of DNMs within recurrently mutated elements, which might have highlighted disruption of specific binding sites or functional sub-regions within each element.

**Figure 3.**
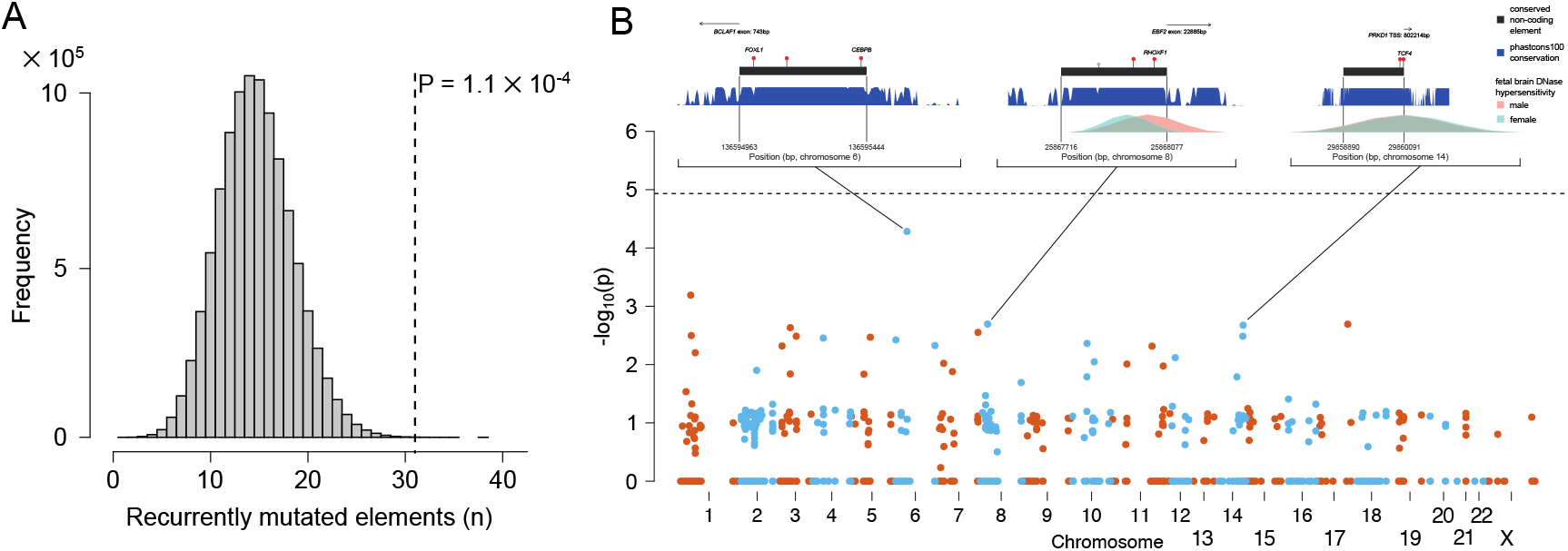
Enrichment of recurrently mutated non-coding elements. (a) Distribution of number of recurrently mutated fetal brain active non-coding elements expected under the null model. Vertical line indicates observed count. (b) Enrichment test of individual non-coding elements. No individual element was significant at a genome-wide threshold of P < 1.9 × 10^−5^ (Bonferroni correction for testing 2,613 fetal brain active elements). Inset plots for three elements show the nearest exon or transcription start site, location of DNMs (red markers), location of rare variants in unaffected parents (grey markers), evolutionary conservation (blue, higher indicates more conserved), and fetal brain DNase hypersensitivity (male in pink, female in blue).

Increased power to detect locus-specific enrichments of DNMs could, in principle, be gained from aggregating DNMs across elements regulating the same target gene, analogous to the aggregation of DNMs in distinct exons of the same protein-coding gene. However, as described above, gene target prediction is currently lacking in both coverage and accuracy. CNEs have been shown to cluster together within the genome^32^, largely due to multiple regulatory elements acting on the same target gene. Moreover, CNEs are strongly enriched around developmentally important genes^32^. Therefore we clustered CNEs spatially to identify groups of CNEs likely to be regulating the same target gene, while remaining agnostic as to the identity of that gene. We applied hierarchical clustering on the 2,613 fetal brain active CNEs to identify 356 clusters of two or more fetal brain active CNEs (Methods). We found an excess of recurrently mutated clusters, defined as two or more elements with at least one DNM in each element, (11 observed, 6 expected, p = 0.016). As with individual elements, we did not find any element clusters with a significant excess of DNMs at a genome-wide significance threshold (Table S1). We compared phenotypic similarity between probands with DNMs in the same CNE or CNE-cluster and randomly sampled probands (Methods) and found probands with DNMs in the same element or cluster to be more phenotypically similar than expected by chance (p = 0.013, one-sided KS-test comparing observed p-values to expected uniform distribution)

We used chromHMM^26^ to assign the recurrently mutated CNEs to a predicted chromatin state. We observed the greatest excess of DNMs in CNEs predicted to be enhancers (N=9) or strongly/weakly transcribed (N=8). Five of the eight transcribed recurrently mutated elements fall in close proximity to exons, but are not in protein-coding transcripts and show evidence for involvement in alternative splicing or induction of nonsense-mediated decay (*BCLAF1, SRRT, SLC10A7*, and MKNK1) or as a 3’ UTR (*CELF1*) (Figure S10). The remaining three transcribed recurrently mutated elements overlap DHS peaks and may be weakly transcribed enhancers or long non-coding RNAs. The full set of recurrently mutated elements is described in Table 1 and the location of DNMs, population variation, and additional annotations are shown in Figure S11.

**Table 1.** Genomic coordinates of the 31 recurrently mutated fetal brain active CNEs and conserved enhancers. Annotated with the nearest gene and any gene interaction reported by Hi-C data in the fetal brain. P-value is the probability of observing at least as many as the reported number of DNMs in the 6,239 exome-negative probands under the null mutation model.

| Region | Class | DNMs | p-value | Nearest Gene | Hi-C Gene Interactions |
| --- | --- | --- | --- | --- | --- |
| chr6:136594963-136595444 | Conserved | 3 | 5.21E-05 | BCLAF1 | AHI1,MAP7 |
| chr1:47040703-47041712 | Conserved | 3 | 0.000644352 | MKNK1 | NONE |
| chr8:25867716-25868077 | Conserved | 2 | 0.002022041 | EBF2 | EBF2,DOCK5 |
| chr17:2095049-2095410 | Conserved | 2 | 0.002027063 | SMG6 | NONE |
| chr14:57553276-57553757 | Conserved | 2 | 0.002115533 | EXOC5 | NONE |
| chr3:71277252-71279016 | Enhancer | 3 | 0.002326107 | FOXP1 | NONE |
| chr7:114999315-114999916 | Conserved | 2 | 0.002809456 | MDFIC | FOXP2 |
| chr1:87795132-87796797 | Enhancer | 3 | 0.003179772 | LMO4 | NONE |
| chr3:136760251-136760852 | Conserved | 2 | 0.003260206 | IL20RB | MSL2,PPP2R3A,STAG1,CLDN18 |
| chr14:37558933-37559534 | Conserved | 2 | 0.003269219 | SLC25A21 | TTC6 |
| chr5:163987267-163987868 | Conserved | 2 | 0.003394978 | MAT2B | NONE |
| chr4:147215258-147215935 | Conserved | 2 | 0.003512212 | SLC10A7 | NONE |
| chr6:14501358-14501959 | Conserved | 2 | 0.003777185 | CD83 | NONE |
| chr10:131699490-131700091 | Conserved | 2 | 0.004335706 | EBF3 | NONE |
| chr6:91341961-91342682 | Conserved | 2 | 0.004717106 | MAP3K7 | NONE |
| chr3:19028185-19028906 | Conserved | 2 | 0.00479714 | KCNH8 | NONE |
| chr11:8310678-8311279 | Conserved | 2 | 0.004835448 | LMO1 | NONE |
| chr1:90847520-90848241 | Conserved | 2 | 0.006271403 | BARHL2 | NONE |
| chr12:17033589-17034430 | Conserved | 2 | 0.00760719 | LMO3 | NONE |
| chr10:103245609-103246330 | Conserved | 2 | 0.008960215 | BTRC | LBX1 |
| chr7:100480136-100480803 | Conserved | 2 | 0.009519587 | SRRT | MUC17 |
| chr11:20297786-20298693 | Conserved | 2 | 0.009776495 | HTATIP2 | NONE |
| chr11:47487425-47488506 | Conserved | 2 | 0.010574131 | CELF1 | MTCH2 |
| chr2:60077201-60078311 | Conserved | 2 | 0.012569209 | BCL11A | NONE |
| chr7:13506147-13507336 | Enhancer | 2 | 0.013216064 | ETV1 | NONE |
| chr3:180461765-180462934 | Enhancer | 2 | 0.014485741 | CCDC39 | NONE |

## Estimating the genome-wide burden of DNMs in fetal brain active elements

The absence of individual non-coding elements with a genome-wide significant enrichment of DNMs allowed us to place an upper bound on the proportion of sites and elements at which DNMs are pathogenic. Approximately 8% of DNMs in protein-coding regions result in a protein-truncating mutation^25,33^. Unlike protein-coding regions, CNEs lack annotation to identify putative pathogenic mutations and are smaller (median 600bp). Downsampling gene length to 600bp and masking annotation of protein consequence, results in an 80% drop in empirical power for the 94 genes passing the genome-wide significance threshold in McRae et. al, 2017 (Figure S12a). This empirical estimate is close to the simulated estimate (Figure S12b). As we did not discover any genome-wide significant CNEs, the proportion of DNMs in CNEs that are pathogenic and highly penetrant must be substantially lower than 8%. Figure 4a depicts the likelihood of observing zero genome-wide significant non-coding elements given the number of fetal brain active CNEs (out of 2,613) in which DNMs can cause neurodevelopmental disorders and the proportion of mutations in those elements that are pathogenic. This analysis suggests that very few DNMs in these elements (likely <2%) are highly penetrant for dominant neurodevelopmental disorders.

**Figure 4.**
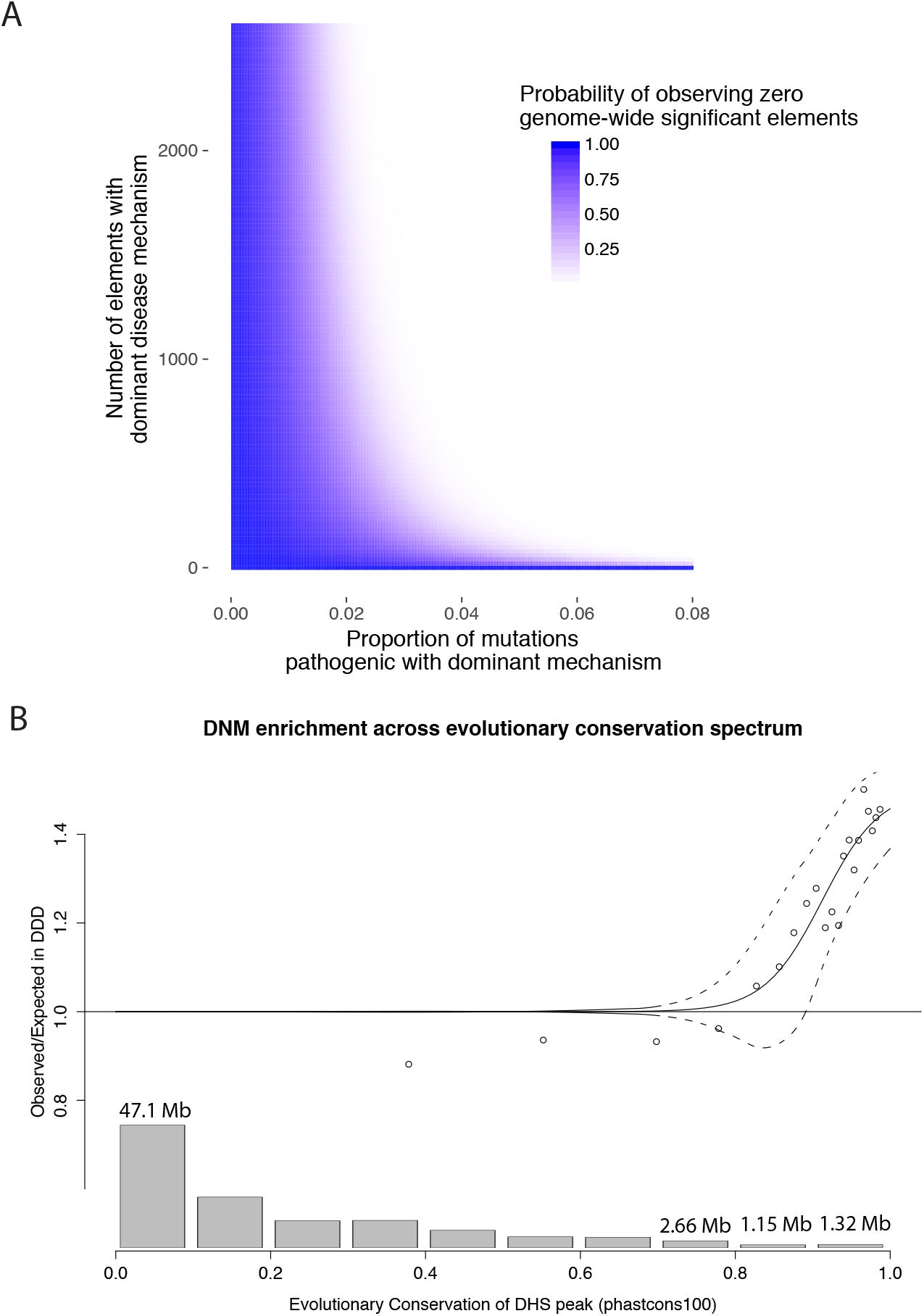
Modelling the proportion of DNMs in non-coding elements that are likely to be highly penetrant for dominant neurodevelopmental disorders. (a) Based on our observation of zero noncoding elements at genome-wide significance in more than 6,000 exome-negative probands, very few sites within these elements (<2%) are likely to contribute to developmental disorders through a highly penetrant dominant mechanism. (b) Logistic regression used to model the genome-wide contribution of dominant-acting DNMs in fetal brain DNase hypersensitive sites in non-coding elements as a function of level of evolutionary conservation. Dashed lines indicate the upper and lower 95% CI. Barplot across the bottom shows the fetal brain active DHS peaks genome-wide at a given level of evolutionary conservation (in megabases).

Our survey of the non-coding genome is biased towards elements with a high degree of evolutionary conservation, but also includes fetal brain active elements with lower levels of evolutionary conservation. To extrapolate the excess of DNMs we observed in the targeted fetal-brain active elements genome-wide, we fitted the enrichment of DNMs as a function of evolutionary conservation using logistic regression (Methods). Factoring in the distribution of evolutionary conservation of all fetal brain DHS peaks genome-wide, we predicted a genome-wide excess of 88 DNMs (95% CI: 48-140) across the 4,915 exome-negative probands with neurodevelopmental disorders, corresponding to 1.0% – 2.8% of unsolved cases (Figure 4b).

## Discussion

In summary, we have demonstrated that severe neurodevelopmental disorders can be caused by highly penetrant *de novo* mutations in regulatory elements. The majority of these regulatory elements likely act as enhancers. This significant excess of DNMs is only observed in the most highly evolutionary conserved elements that are active in the fetal brain. These elements also exhibit substantial selective constraint within human populations. We observed a 1.3-fold excess of DNMs within DHS peaks within these regulatory elements, suggesting that only a minority of such DNMs are pathogenic. Moreover, our modelling suggests that the proportion of all possible mutations within these elements that are likely to cause dominant neurodevelopmental disorders is very small (<2%). As a consequence, this class of pathogenic non-coding DNMs is only likely to account for a small proportion (<5%) of ‘exome-negative’ individuals.

The nature of our study design focuses on highly conserved elements and fetal-brain active elements and is relatively uninformative with respect to pathogenic ‘gain-of-function’ DNMs within elements that show no wildtype activity in fetal brain, and are not highly evolutionarily conserved. Recently, targeted sequencing in 218 consanguineous autism families of ‘human accelerated regions’ (HARs) - non-coding elements that exhibit evolutionary conservation but excessive human-specific divergence - identified a significant 1.43-fold enrichment of rare biallelic variants^34^. We note that the majority of pathogenic DNMs in protein-coding genes causing neurodevelopmental disorders occur in genes that are highly evolutionary conserved, are under strong selective constraint within human populations and exhibit wildtype expression in fetal brain^17,35^. Therefore, the relatively modest excess of DNMs that we observed in this study is highly likely to be an upper bound for other classes of non-coding elements. Consequently, a comprehensive analysis of the contribution of variation within all classes of non-coding elements to neurodevelopmental disorders is likely to require whole genome sequencing (WGS) of many tens of thousands, if not hundreds of thousands of parent-proband trios (Figure S13).

One challenge of interpreting WGS data is the vast universe of different hypotheses that could be tested, and thus how to account appropriately for multiple hypothesis testing. *Turner et al.* recently reported a nominally significant enrichment (p = 0.03) of *de novo* SNVs and private copy number variants in fetal brain DHS or at sites with PhyloP conservation score of >4, within 50kb of known autism-associated genes in whole genome sequences from 53 individuals with autism^36^. Caution should be exercised in interpreting findings based on: small sample sizes relative to that required for well-powered analyses (as discussed above), especially in cohorts with lower mutational burden (e.g. within protein-coding genes) – and therefore lower statistical power - compared to the cohort studied here, and analyses requiring multiple, arbitrary, levels of variant stratification (e.g. gene set, genomic proximity threshold, and conservation score). WGS-based analyses need to explicitly articulate and account for all tested hypotheses.

Our analyses were limited to SNVs as current models for indel mutation processes are too inaccurate to allow robust assessment of mutational excess. In addition, our analyses also highlight an urgent need for improved tools to stratify more and less damaging variants within non-coding elements, and more accurate and comprehensive annotation of gene targets for individual regulatory elements. These improved mutational models and functionally-relevant annotations will dramatically increase power to detect highly-penetrant disease-associated non-coding variation, for example, increasing power more than ten-fold from 5% to 83% in 40,000 trios (Figure S13). Functional characterisation of increasing numbers of robustly-associated, highly-penetrant, regulatory variants in cellular and animal models will be critical in moving us from a descriptive to a more predictive understanding of non-coding variation in the human genome, as well as elucidating their underlying pathophysiological mechanisms.

## Acknowledgements

We thank the families for their participation and patience. We are grateful to the Exome Aggregation Consortium for making their data and code available. We would like to thank Sebastian Gerety, Greg Elgar, Stein Aerts, and Dmitry Svetlichnyy for helpful discussions. We thank Hugues Roest Crollius and Lambert Moyon for help on gene target prediction. We would like to thank Jonathan Mudge and Adam Frankish for their help in annotation of conserved non-coding elements. We thank the Sanger HGI and DNA pipelines teams for their support in generating and processing the data. The DDD study presents independent research commissioned by the Health Innovation Challenge Fund (grant HICF-1009-003), a parallel funding partnership between the Wellcome Trust and the UK Department of Health, and the Wellcome Trust Sanger Institute (grant WT098051). The views expressed in this publication are those of the author(s) and not necessarily those of the Wellcome Trust or the UK Department of Health. The study has UK Research Ethics Committee approval (10/H0305/83, granted by the Cambridge South Research Ethics Committee and GEN/284/12, granted by the Republic of Ireland Research Ethics Committee). D.R.F. is funded through an MRC Human Genetics Unit program grant to the University of Edinburgh.

## References

1 Hindorff, L. a. et al. Potential etiologic and functional implications of genome-wide association loci for human diseases and traits. Proceedings of the National Academy of Sciences of the United States of America 106, 9362–9367, doi:10.1073/pnas.0903103106 (2009).

2 Maurano, M. T. et al. Systematic Localization of Common Disease-Associated Variation in Regulatory DNA. Science 337, 1190–1195, doi:10.1126/science.1222794 (2012).

3 Mathelier, A., Shi, W. & Wasserman, W. W. Identification of altered cis-regulatory elements in human disease. Trends in Genetics 31, 67–76, doi:10.1016/j.tig.2014.12.003 (2015).

4 Spielmann, M. & Mundlos, S. Looking beyond the genes: the role of noncoding variants in human disease. 25, 157–165, doi:10.1093/hmg/ddw205 (2016).

5 Weedon, M. N. et al. Recessive mutations in a distal PTF1A enhancer cause isolated pancreatic agenesis. Nature genetics 46, 61–64, doi:10.1038/ng.2826 (2014).

6 Jeong, Y. et al. Regulation of a remote Shh forebrain enhancer by the Six3 homeoprotein. Nature genetics 40, 1348–1353, doi:10.1038/ng.230 (2008).

7 Benko, S. et al. Disruption of a long distance regulatory region upstream of SOX9 in isolated disorders of sex development. Journal of medical genetics 48, 825–830, doi:10.1136/jmedgenet-2011-100255 (2011).

8 Bhatia, S. et al. Disruption of autoregulatory feedback by a mutation in a remote, ultraconserved PAX6 enhancer causes aniridia. American Journal of Human Genetics 93, 1126–1134, doi:10.1016/j.ajhg.2013.10.028 (2013).

9 Weedon, M. N. et al. Recessive mutations in a distal PTF1A enhancer cause isolated pancreatic agenesis. Nature genetics 46, 61–64, doi:10.1038/ng.2826 (2014).

10 Lettice, L. A. et al. A long-range Shh enhancer regulates expression in the developing limb and fin and is associated with preaxial polydactyly. Human Molecular Genetics 12, 1725–1735, doi:10.1093/hmg/ddg180 (2003).

11 Hill, R. E., Lettice, L. A. & Hill, R. E. Alterations to the remote control of Shh gene expression cause congenital abnormalities. (2013).

12 Noonan, J. P. & McCallion, A. S. Genomics of Long-Range Regulatory Elements. Annual Review of Genomics and Human Genetics 11, 1–23, doi:10.1146/annurev-genom-082509-141651 (2010).

13 Naville, M. et al. Long-range evolutionary constraints reveal cis-regulatory interactions on the human X chromosome. Nature Communications 6, 6904, doi:10.1038/ncomms7904 (2015).

14 Whalen, S., Truty, R. M. & Pollard, K. S. Enhancer-promoter interactions are encoded by complex genomic signatures on looping chromatin. Nature genetics 48, 488–496, doi:10.1038/ng.3539 (2016).

15 Köhler, S. et al. The Human Phenotype Ontology in 2017. Nucleic acids research 45, D865–D876, doi:10.1093/nar/gkw1039 (2017).

16 Wright, C. F. et al. Genetic diagnosis of developmental disorders in the DDD study: a scalable analysis of genome-wide research data. The Lancet 385, 1305–1314, doi:10.1016/S0140-6736(14)61705-0 (2015).

17 Deciphering Developmental Disorders, S. Prevalence and architecture of de novo mutations in developmental disorders. Nature, doi:10.1038/nature21062 (2017).

18 Siepel, A. et al. Evolutionarily conserved elements in vertebrate, insect, worm, and yeast genomes. Genome Research 15, 1034–1050, doi:10.1101/gr.3715005 (2005).

19 Visel, A., Minovitsky, S., Dubchak, I. & Pennacchio, L. A. VISTA Enhancer Browser - A database of tissue-specific human enhancers. Nucleic Acids Research 35, 88–92, doi:10.1093/nar/gkl822 (2007).

20 May, D. et al. Large-scale discovery of enhancers from human heart tissue. Nature genetics 44, 89–93, doi:10.1038/ng.1006 (2012).

21 Lek, M. et al. Analysis of protein-coding genetic variation in 60,706 humans. Nature Publishing Group 536, 285–291, doi:10.1038/nature19057 (2016).

22 Kircher, M. et al. A general framework for estimating the relative pathogenicity of human genetic variants. Nature genetics 46, 310–315, doi:10.1038/ng.2892 (2014).

23 Walter, K. et al. The UK10K project identifies rare variants in health and disease. Nature 526, 82–90, doi:10.1038/nature14962 (2015).

24 Consortium, R. E. et al. Integrative analysis of 111 reference human epigenomes. Nature 518, 317–330, doi:10.1038/nature14248 (2015).

25 Samocha, K. E. et al. A framework for the interpretation of de novo mutation in human disease. Nature genetics 46, 944–950, doi:10.1038/ng.3050 (2014).

26 Ernst, J. & Kellis, M. ChromHMM: automating chromatin-state discovery and characterization. Nature methods 9, 215–216, doi:10.1038/nmeth.1906 (2012).

27 Pollard, K. S., Hubisz, M. J., Rosenbloom, K. R. & Siepel, A. Detection of nonneutral substitution rates on mammalian phylogenies. Genome Research 20, 110–121, doi:10.1101/gr.097857.109 (2010).

28 Louis, a., Nguyen, N. T. T., Muffato, M. & Roest Crollius, H. Genomicus update 2015: KaryoView and MatrixView provide a genome-wide perspective to multispecies comparative genomics. Nucleic Acids Research 43, D682–D689, doi:10.1093/nar/gku1112 (2014).

29 Shooshtari, P., Huang, H. & Cotsapas, C. Integrative genetic and epigenetic analysis uncovers regulatory mechanisms of autoimmune disease. doi:10.1101/054361 (2016).

30 Won, H. et al. Chromosome conformation elucidates regulatory relationships in developing human brain. Nature Publishing Group 538, 523–527, doi:10.1038/nature19847 (2016).

31 Mathelier, A. et al. JASPAR 2016: A major expansion and update of the open-access database of transcription factor binding profiles. Nucleic Acids Research 44, D110–D115, doi:10.1093/nar/gkv1176 (2016).

32 Sandelin, A. et al. Arrays of ultraconserved non-coding regions span the loci of key developmental genes in vertebrate genomes. BMC genomics 5, 99, doi:10.1186/1471-2164-5-99 (2004).

33 Kryukov, G. V., Pennacchio, L. a. & Sunyaev, S. R. Most rare missense alleles are deleterious in humans: implications for complex disease and association studies. American journal of human genetics 80, 727–739, doi:10.1086/513473 (2007).

34 Doan, R. N. et al. Mutations in Human Accelerated Regions Disrupt Cognition and Social Behavior Article Mutations in Human Accelerated Regions Disrupt Cognition and Social Behavior. Cell 167, 341–354.e312, doi:10.1016/j.cell.2016.08.071 (2016).

35 Purcell, S. M. et al. A polygenic burden of rare disruptive mutations in schizophrenia. Nature 506, 185–190, doi:10.1038/nature12975 (2014).

36 Turner, Tychele N. et al. Genome Sequencing of Autism-Affected Families Reveals Disruption of Putative Noncoding Regulatory DNA. The American Journal of Human Genetics 98, 58–74, doi:10.1016/j.ajhg.2015.11.023 (2016).

37 Harrow, J. et al. GENCODE: the reference human genome annotation for The ENCODE Project. Genome research 22, 1760–1774, doi:10.1101/gr.135350.111 (2012).

38 Gentleman, R. et al. Bioconductor: open software development for computational biology and bioinformatics. Genome Biology 5, R80, doi:10.1186/gb-2004-5-10-r80 (2004).

39 Smedley, D. et al. A Whole-Genome Analysis Framework for Effective Identification of Pathogenic Regulatory Variants in Mendelian Disease. The American Journal of Human Genetics 99, 595–606, doi:10.1016/j.ajhg.2016.07.005 (2016).

40 Shihab, H. a. et al. An integrative approach to predicting the functional effects of non-coding and coding sequence variation. Bioinformatics (Oxford, England) 31, 1536–1543, doi:10.1093/bioinformatics/btv009 (2015).

41 Lawrence, M. et al. Software for computing and annotating genomic ranges. PLoS Comput Biol 9, e1003118, doi:10.1371/journal.pcbi.1003118 (2013).

42 Ramu, A. et al. DeNovoGear: de novo indel and point mutation discovery and phasing. 10, 3–7, doi:10.1038/nmeth.2611 (2013).

43 Akawi, N. et al. Discovery of four recessive developmental disorders using probabilistic genotype and phenotype matching. Nature Publishing Group 47, 1363–1369, doi:10.1038/ng.3410 (2015).

44 Sanyal, A., Lajoie, B. R., Jain, G. & Dekker, J. The long-range interaction landscape of gene promoters. Nature 489, 109–113, doi:10.1038/nature11279 (2012).

45 Consortium, G. T. The Genotype-Tissue Expression (GTEx) project. Nat Genet 45, 580–585, doi:10.1038/ng.2653 (2013).

46 McLeay, R. C. & Bailey, T. L. Motif Enrichment Analysis: a unified framework and an evaluation on ChIP data. BMC bioinformatics 11, 165, doi:10.1186/1471-2105-11-165 (2010).

